# Decoding the Impact of Neighboring Amino Acid on ESI-MS Intensity Output through Deep Learning

**DOI:** 10.1101/2024.02.02.578588

**Authors:** Naim Abdul-Khalek, Reinhard Wimmer, Michael Toft Overgaard, Simon Gregersen Echers

## Abstract

Peptide-level quantification using mass spectrometry (MS) is no trivial task as the physicochemical properties affect both response and detectability. The specific amino acid (AA) sequence affects these properties, however the link from sequence to intensity output remains poorly understood. In this work, we explore combinations of amino acid pairs (i.e., dimer motifs) to determine a potential relationship between the local amino acid environment and MS1 intensity. For this purpose, a deep learning (DL) model, consisting of an encoder-decoder with an attention mechanism, was built. The attention mechanism allowed to identify the most relevant motifs. Specific patterns were consistently observed where a bulky/aromatic and hydrophobic AA followed by a cationic AA as well as consecutive bulky/aromatic and hydrophobic AAs were found important for the MS1 intensity. Correlating attention weights to mean MS1 intensities revealed that some important motifs, particularly containing Trp, His, and Cys, were linked with low responding peptides whereas motifs containing Lys and most bulky hydrophobic AAs were often associated with high responding peptides. Moreover, Asn–Gly was associated with low MS1 response. The model could predict MS1 response with a mean average percentage error of ∼11% and a Pearson correlation coefficient of ∼0.68.

## 1. Introduction

Qualitative and quantitative protein analysis is central in biotechnology, biology, and medicine, but is also gaining headway in new fields such as agriculture and food science [1–3]. For this purpose, mass spectrometry (MS)-based proteomics has undergone tremendous development over the recent decades facilitated by considerable advances in software and instrumentation, allowing for not only qualitative but also quantitative analysis of biological samples[4]. During routine MS analysis, biological samples undergo several processing steps, going from sample preparation, analyte separation (often liquid chromatography, LC), followed by detection in the mass analyzer [5]and finally, the MS outputs are processed with bioinformatic tools. In each step of this workflow, there are multiple factors potentially introducing variability and thereby affecting the results and reproducibility of the analysis [6–12]. This becomes particularly evident when moving from the qualitative to the quantitative domain of MS analysis and downstream application of quantitative data for construction of predictive models using artificial intelligence [13].

On the protein level, quantitative variability observed in MS analysis, is often addressed through a range of operations including imputations and data normalization using certain assumptions [14–16]. Such assumptions may indeed be fair approximations when working with protein level quantification, where multiple datapoints (peptides) are available for each protein but fall short in peptide-centric analysis [17]. Similarly, to other analytes, peptides adhere to the fundamental prerequisite for MS-based quantification that within a certain dynamic range, the relationship between concentration and response can be regarded linear [17]. Under the assumption that physicochemical properties such as chromatographic retention and ionization behavior remains constant between measurements, this allows for relative, ratio-based peptide-level quantification between samples analyzed under the same preparative and analytical conditions. However, such approaches become unfeasible for label-free, in-sample comparison of different peptides due to the large variations in intrinsic ionization properties of individual peptides, which rely on both their amino acid (AA) constituents and their order [18]. While peptide precursor (MS1) intensities from LC-MS/MS analysis have previously been used to approximate peptide-level concentrations [19,20], such approaches will include significant risks of substantially incorrect quantitative estimates. Nevertheless, the ability to deduce peptide quantity by precursor response during label-free LC-MS/MS analysis of highly complex peptide mixtures would be highly relevant for different purposes such as quantitative differentiation of protein isoforms and characterization of protein hydrolysates. Moreover, a better quantitative determination on the peptide level may lead to more accurate determination of absolute, protein-level abundances.

Through both experimental and computational approaches, the global, physicochemical properties of a peptide have been highlighted as highly relevant factors for the behavior of individual peptides during MS analyses [21–27]. Among these properties, hydrophobicity is one of the most relevant features for peptide detectability by seemingly increasing the ionization efficiency of peptides (until certain degree) in Reversed-phase LC–MS [28]. Charge is another highly relevant property, indicating that amino acids (AA) such as arginine (Arg) or lysine (Lys) in a peptide sequence will increase the likelihood of generating a positively charged through side chain protonation under the acidic pH conditions employed is LC-ESI-MS [29]. Moreover, structural propensities have been identified to be relevant as well. It has been argued that peptides with known and stable α-helical and β-sheet structures in solution might have negative effect on the intensity output of the MS analysis [21].

Since physicochemical properties of peptides greatly affect the ionization potential and thus response during MS analysis, gaining a better understanding of their influence is a key step for not only designing experiment that will yield better result, but also for continued development of current computational technologies. In previous work [27], we investigated the relevancy of each AA on the MS1 response, but results indicated that single AA occurrence was not sufficient to describe the response variability in a generalizable manner. The logical extension of this is to consider if specific combinations of AAs have a synergistic or antagonistic effect and may be regarded better descriptors of the MS1 response on a more general plane. Thus, in this work we present a deep learning (DL) model with an attention mechanism and employ it to identify and explore the possible impact of AA pairs (dimer motifs). Through this, we investigate if certain combinations affect the response more than others and if considering peptides as sequences of dimers may lead to an improved predictability of the MS1 response.

## 2. RESULTS AND DISCUSSION

The dataset was initially inspected and filtered to remove redundancies, errors, and reduce noise, whereafter the cumulative AA frequency was explored. The most abundant AAs are leucine (Leu), serine (Ser) and glutamic acid (Glu), while the least abundant AA are tryptophan (Trp), methionine (Met), and cysteine (Cys) (Fig. 1A). Considering the number of peptides containing any specific AA (Fig. 1B), there are some clear differences compared to the cumulative AA distribution. The most noticeable difference is that arginine (Arg) is observed in the second highest number of peptides (ranked 7 in the cumulative distribution) but also an increase in the prevalence of lysine (Lys) is apparent (from rank 9 in the cumulative distribution to rank 5). This can be attributed to the fact that the dataset contains only peptides identified by *in silico* tryptic digestion, thus the vast majority of peptides will contain Arg or Lys at the C-terminus.

**Figure 1.**
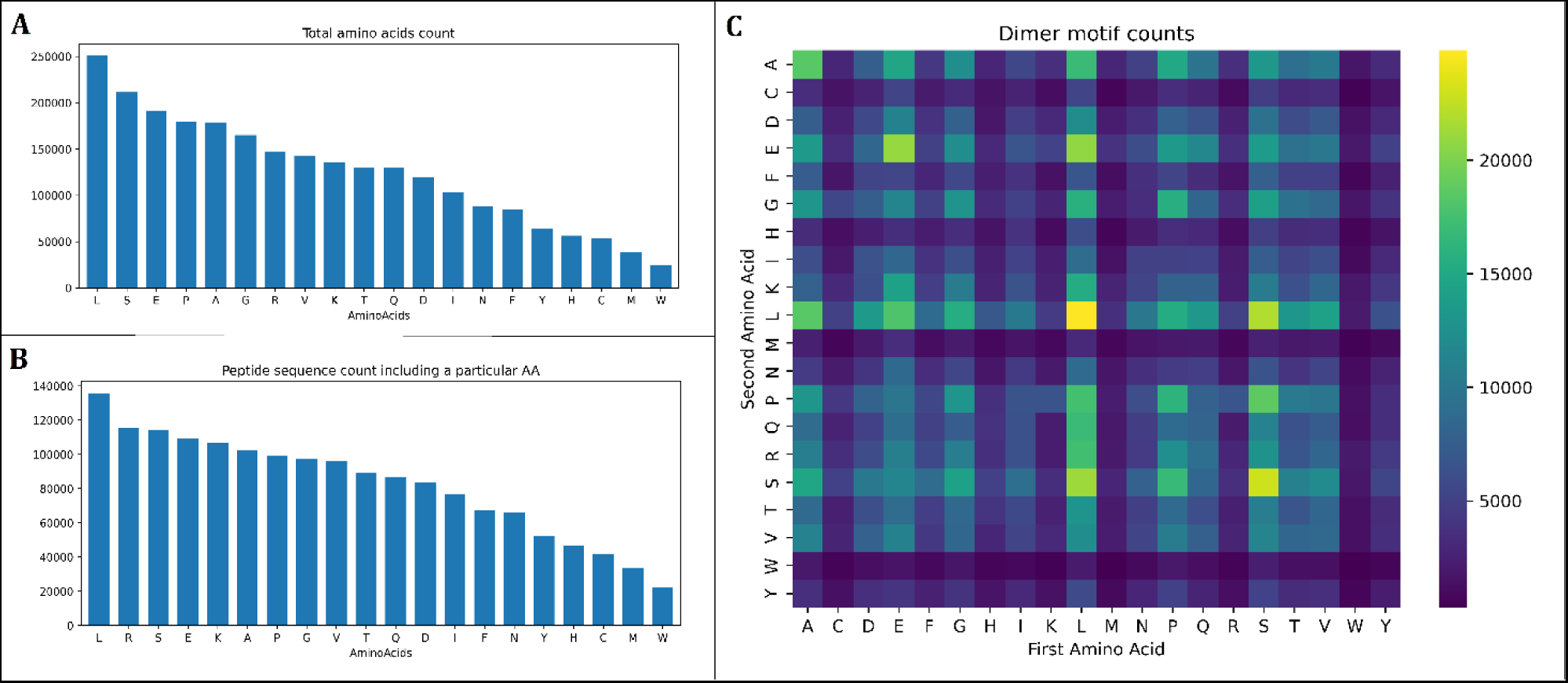
Graphical representation of AA frequencies in filtered dataset. A) Histogram of total AA count (total: 2,494,871 counts). B) Histogram of peptide sequence count including a particular AA (total: 1,642,595 counts). C) Heatmap of dimer motif counts (total: 2,315,649 counts).

Looking into the frequency of dimer motifs (Fig. 1C, Table S2), a high level of similarity with the total AA distribution (Fig. 1A) was intuitively observed, showing Leu, Ser, Glu, and combinations hereof as the most frequent. Moreover, some motifs of two identical AAs (the diagonal, i.e., Leu-Leu, Ser-Ser, Glu-Glu, but also Ala-Ala) were particularly frequent (Fig. 1C). The frequency of individual AAs and dimer motifs is important for evaluating their impact on MS1 response. For instance, if a particular motif is found with very low frequency, a small subset of datapoints may be recognized by the model as important, but due to the low sample size of this particular feature, the statistical uncertainty may be too large to reach any meaningful conclusion and introduce bias in the model. In a recent study, we investigated the effect of individual AAs on the peptide-level MS1 response using the same dataset [27]. Here, we identified Trp, Leu, Arg, and Lys, among others, to be important for governing the MS1 response. Curiously, these AAs were among both the most and least frequent considering both cumulative and peptide counts (Fig. 1A, 1B). This could indicate potential bias, inflicted by the overrepresentation of e.g. Leu and underrepresentation of e.g. Trp in nature [30], which is reflected in the dataset (Fig. 1A). While some motifs are almost two orders of magnitude overrepresented compared to others (e.g. Leu-Leu (24819) and Trp-Trp (306), Table S2), the substantial number of datapoints for each AA should alleviate this concern and the findings are still considered valid.

### 2.1. Representative Models

Different models were obtained using various sets of hyperparameters during training and optimization. Those models have similar predictive performance. However, the models displayed varying discrepancies with regards to the attention weights for the AA dimers. Three patterns were consistently observed by the attention mechanism, and representative models (RMs) of these patterns are presented below.

#### 2.1.1. Representative model 1: Bulky hydrophobic/aromatic AAs followed by a positively charged AA

The attention weights of the first representative model (RM1) point to a combination of a bulky hydrophobic/aromatic AA followed by a positively charged AA being particularly relevant for the prediction (Fig. 2A). In the first position, phenylalanine (Phe), Trp, Leu, and Isoleucine (Ile) received the highest attention and in the second position, Arg appeared of higher importance than Lys, while histidine (His) received substantially lower attention (Fig. 2B, Table S3). The top 4 dimers were combinations of Phe, Ile, Leu, and Trp in the first position and Arg in the second position (Fig. 2A). Phe and Trp are bulky, aromatic, and hydrophobic AAs while Leu and Ile are regarded as bulky hydrophobic AAs. Interestingly, the combination Tyr-Arg and methionine (Met)-Arg receives lower attention than valine (Val)-Arg, despite Val being smaller than Tyr and Met. This could be ascribed to the presence of non-hydrocarbon moieties (-OH and -S- in Tyr and Met, respectively), which increase side chain polarity. While bulkiness and aromaticity appear to be important, the properties also appear to require a certain balance with side-chain hydrophobicity. Moreover, Met is prone to oxidation under physiological conditions [31], during sample preparation [32], and in the gas-phase following ionization [33]. The product of oxidation, methionine sulfoxide, has been reported to be reactive [31] but also susceptible to selective fragmentation within the side chain – particularly in the vicinity of Arg [34]. The interaction between Tyr and Arg has previously been reported as the most abundantly observed cation-π interaction in proteins [35], and have also been linked with formation of non-covalent complexes [36] and side-chain loss [37] in the gas-phase. These factors could lead to a generally lower response than expected for the unmodified precursor ion.

**Figure 2.**
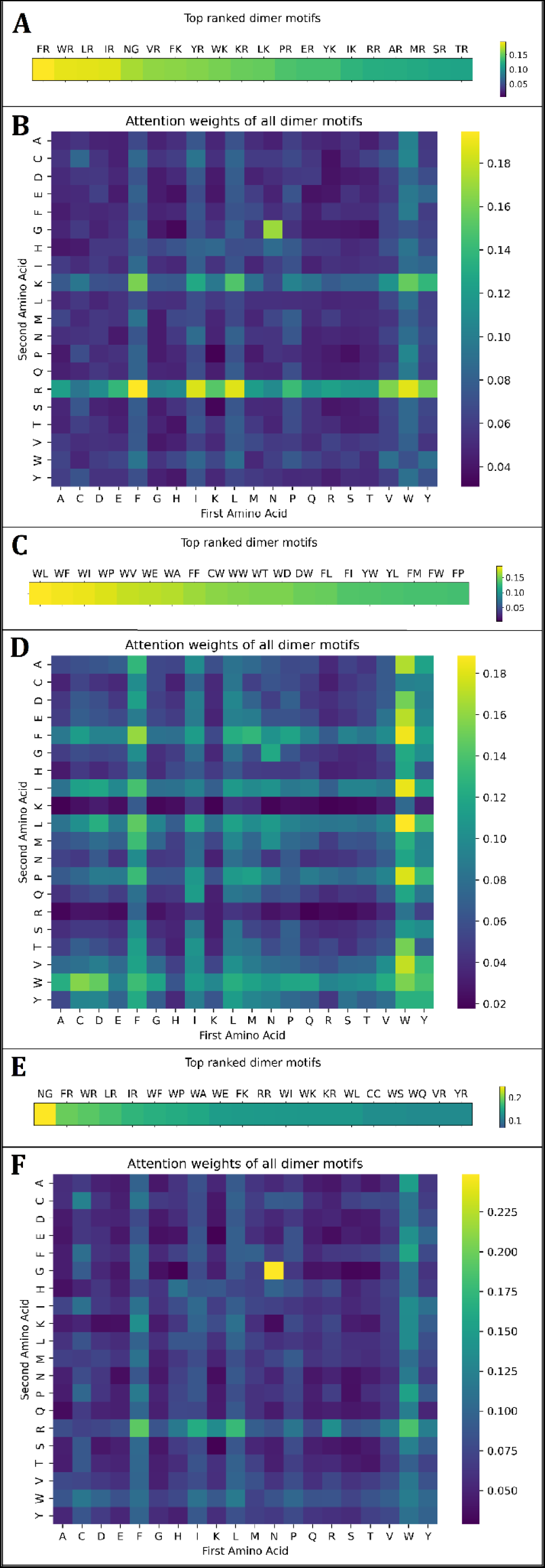
Heatmaps of attention weights for representative models (RMs) 1, 2, and 3. A) Top 20 dimer motif weights for RM 1. B) Matrix of all dimer motif weights for RM 1. C) Top 20 dimer motif weights for RM 2. D) Matrix of all dimer motif weights for RM 2. E) Top 20 dimer motif weights for RM 3. F) Matrix of all dimer motif weights for RM 3.

Notably, if the order of the dimers is inverted (charged–hydrophobic/aromatic), the attention weights are much lower. However, since a tryptic dataset was used and trypsin cuts C-terminally of Arg and Lys, these AAs are predominantly found in the second position of the dimer (Fig. 1C, Table S2) and frequently at the C-terminus of the peptide.

#### 2.1.2. Representative model 2: Aromatic AAs followed by a hydrophobic/aromatic AA

In this second representative model (RM2), attention weights mainly focused on AA combinations with high aromaticity, hydrophobicity, and size (Fig. 2C, 2D, Table S4). In the first position, Trp, Phe, and Tyr (and to a lesser extent Leu and Ile) were the most relevant while in the second position, Trp, Phe, Leu, Ile, and Tyr received the most attention (Fig. 2D). Notably, proline (Pro) also received noteworthy attention in the second position. In this model, Trp generally shows higher relevance than all other AAs in the first position. Moreover, Tyr obtained higher weights in RM 2 than in RM 1. Since positively charged AAs were not considered significant by RM 2, this suggests a possible interaction between its hydroxyl group and the protonated side chain of the adjacent Arg/Lys. This interaction is devoid when Tyr is followed by a non-charged AA, explaining why Tyr becomes more important and comparable to the other aromatic Aas in RM2. Thus, it seems that an interaction between Tyr and an adjacent Arg/Lys, reduce the relevance of Tyr for predicting MS1 response.

As found in RM 1, the order of AAs also appeared important (Fig. 2D), but to a lesser degree (Fig. 2B). RM 2 generally seemed to focus greatly on hydrophobic dipeptide motifs. In previous work, hydrophobicity was also found to be an important feature when only considering single AA occurrence [27]. This observation seems to extend to dimer motifs, thereby adding to the hypothesis that overall and/or local hydrophobicity has a significant impact on MS1 response for peptides. The hypothesis is further corroborated by previous studies, where a positive correlation between ionization efficiency and retention times was empirically demonstrated. [38–40]. Nevertheless, hydrophobic stretches have also been linked with difficulty of synthesis [41], which would result in deletion isoforms of the intended peptide, and thus lower abundance and ultimately response in an intended equimolar synthetic peptide library. Ultimately, hydrophobicity must still be regarded as only one piece of the puzzle, as other studies demonstrated that the correlation between hydrophobicity and ionization efficiency is not linear [26].

#### 2.1.3. Representative model 3: Asn-Gly

RM 3 appeared to overall combine the attentions obtained in RMs 1 and 2. This model highlight dimer motifs with a combination of hydrophobic/aromatic and charged AAs as well as combinations of only hydrophobic/aromatic AAs (Fig. 2F, Table S5). Similarly, to RMs 1 and 2, Trp appears to be of substantial significance for the predictions, pointing to the general importance of this AA, as previously described [27]. However, this model is particularly interesting as the highest attention weight was assigned to the dimer consisting of asparagine and glycine (Asn–Gly) (Fig. 2E). The Asn-Gly motif also received considerable attention in RM 1 (Fig. 2B), where it obtained the fifth highest attention (Fig. 2A) but deviate from the general patterns observed in RMs 1 and 2. Asn is classified as a polar AA with an uncharged side chain while Gly is a very small AA considered as a special case. As no other combination of similar constitution was assigned high attention weights, the motif is, at first glance, not intuitive. Previous studies have reported that peptides containing Asn-Gly typically results in deamidation to an extent of around 70% to 80% [42] but that deamidation is also affected by higher order structure in polypeptides and proteins in solution [43]. In deamidation of Asn, the residue is converted to Asp/βAsp, which may be further modified by aspartimide and diketopiperazine formation [44]. Both deamidation and aspartimide formation is particularly pronounced for Asn-Gly [42,45] and much less pronouced for the inverted motif, Gly-Asn [46]. The highly specific and efficient deamidation of the Asn-Gly motif also explains why the inverted motif did not receive considerable attention (Fig. 2F). An Asn deamidation, and a potential subsequent aspartimide formation, would directly lead to lower precursor abundance unless the modification is explicitly included in the data analysis. While deamidation rate is affected by peptide structure, deamidation may also itself influence peptide structure [47]. Structural change may then again impact MS1 response, as solution- and gas-phase peptide structure was previously identified as an important contributor to MS1 response [27]. Furthermore, inclusion of Asn-Gly is also frequently linked with difficult sequences during peptide synthesis due to aspartimide formation [45,48,49]. As with difficult peptides containing hydrophobic stretches, aspartimide formation will lead to a reduced concentration of the desired peptide, which inevitably will have an effect in the MS1 intensity output. Ultimately, Asn-Gly may negatively affect the MS1 response by reducing precursor abundance through different mechanisms related to synthesizability, stability, and gas-phase reactions. Our model cannot, based on the current data, explicitly differentiate these factors, warranting for further investigation.

### 2.2. Are important dimers reflected directly in the data?

The three RMs all identified several highly relevant dimer motifs, albeit the attention mechanism do not seem to indicate whether the presence of a particular motif has a positive or negative contribution to the MS1 response. Consequently, we investigated the dimer motif-specific intensity distribution (i.e., all peptides containing a specific dimer) within the dataset to determine if these motifs generally stood out. A substantial number of dimer motifs identified by the RMs as relevant were also directly reflected in the intensity data (Fig. 3). Overall, dimers containing Leu, Lys, Ile, Phe, Tyr, Asp, Glu, Val, and Ala generally appeared to have a higher mean MS1 response regardless of position, whereas peptides containing Cys, Trp, and His generally had a lower response. These findings also correlate quite well with previous work on importance of single AA occurrence, where many of these AAs were also identified as important using a similar DL model architecture [27]. Moreover, plotting mean intensities also shed further light on the why the models may have identified certain features as important.

**Figure 3.**
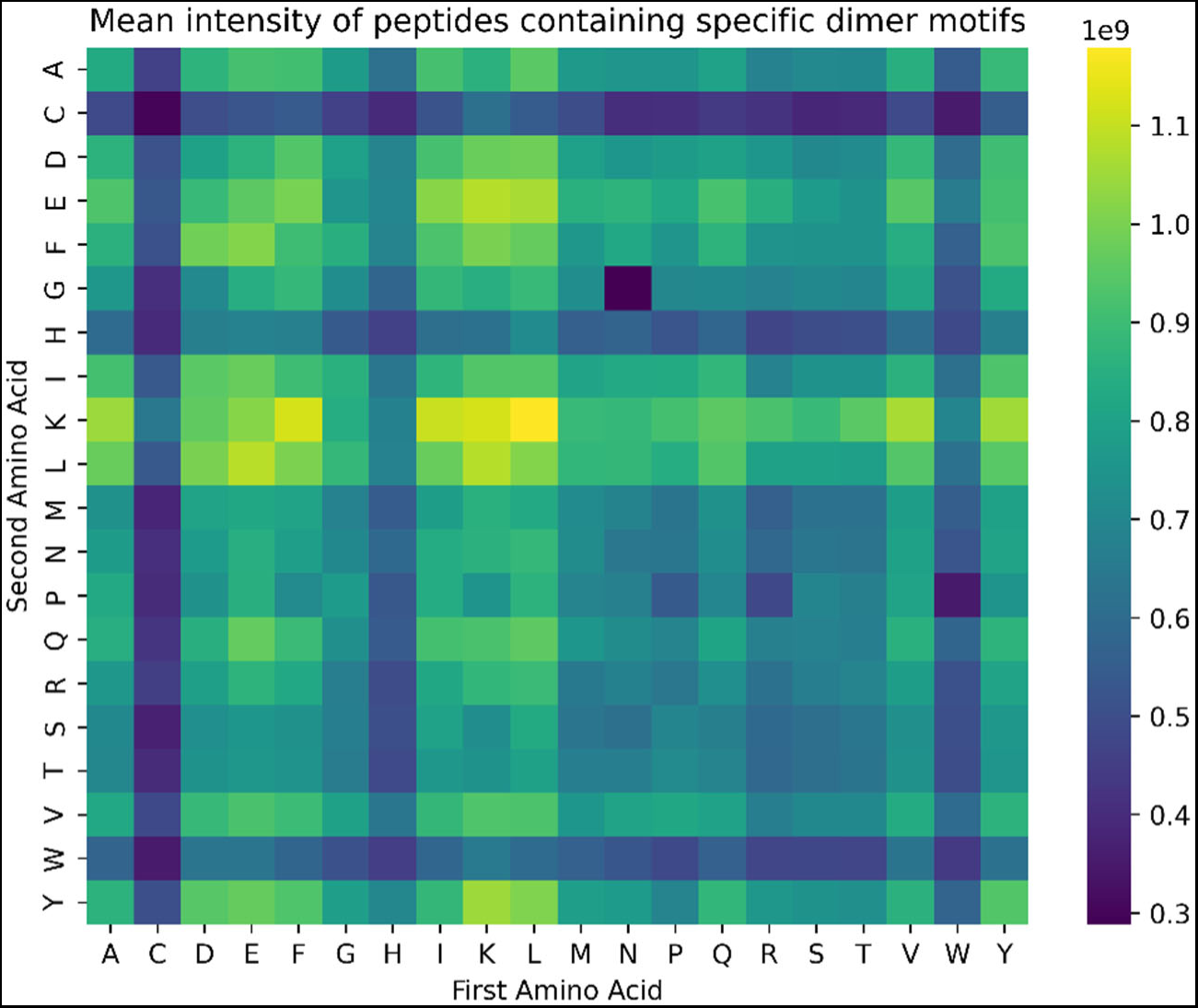
Heatmap representation of mean MS1 intensity for peptides containing specific dimer motifs within the filtered dataset. The plotted intensity is the global mean of any peptide containing a specific dimer motif at any position within the sequence.

Where RM 1 generally attributed dimers with Arg in the second position (Xxx-Arg) as more important than Xxx-Lys motifs (Fig. 2B), intensity data indicate that Xxx-Lys motifs provide higher responses. As both motifs have comparable occurrence in the dataset (Fig. 1C, Table S2), this indicates that RM 1 may be considering the seemingly negative impact of Arg compared to Lys on the MS1 response. This may be attributed to Arg being susceptible to lactamisation during solid-phase synthesis [49]. Another possible reason is that Arg, similarly to the low responders Trp and His, contain higher order (secondar and/or tertiary) side-chain amines, as primary amines have been reported to possess higher threshold energies and a higher probability to survive the ion extraction process compared to secondary amines [50]. This finding could imply that the use of a more specific protease such as LysC could be beneficial compared to trypsin in bottom-up proteomics. However, this would require Lys content and distribution to be compatible with a peptide size-range commonly employed in such analysis.

RM2 focused greatly on aromatic and bulky hydrophobic AAs with particular focus on Trp (Fig. 2D). Interestingly, this particular focus on Trp appears to reflect the overall negative impact that Trp seems to have on the MS1 response regardless of being first or second in the dimer motif, while the remaining aromatic and bulky hydrophobic AAs seem to have a generally positive influence (Fig. 3). This is in contrast to our previous work, where we hypothesized that the importance of Trp was linked with its ability to stabilize the precursor ion through the indole moiety, thus having a positive contribution to the MS1 response [27]. ESI is recognized for accelerating droplet-based reactions [51,52], and Trp has been reported to be labile to both single and double oxidation during sample preparation [53] and electrospray ionization [54,55]. As oxidized Trp has furthermore been reported to rapidly undergo electrochemical modification [56] and cleavage [57] reactions in ESI, this may together explain why Trp-containing peptides generally have lower response in modification-restricted analysis.

RM 3 (Fig. 2F), as well as RM 1 (Fig. 2B), highlighted that the Asn-Gly motif was highly relevant to describe the MS1 response, and intensity data clearly show that this particular dimer motif has a substantial negative effect on the MS1 response. This observation fits well with the motif being particularly susceptible to deamidation and aspartimide formation, thus lowering concentration, and ultimately MS1 response of the unmodified precursor ion, as discussed above. A similar observation of low mean response is found for Trp-Pro (Fig. 3). While this dimer motif did not receive as high attention in the RMs as Ans-Gly, it was identified as important with the fourth highest attention weight in RM 2 (Fig. 2C) and seventh highest weight in RM 3 (Fig. 2E).

Interestingly, the RMs appear to focus more on negative contributors to the MS1 response than dimer motifs with a high mean response. The Leu-Lys motif displayed the highest mean response (Fig. 3) but did not stand out in RM 2 and RM 3, while it ranks 11^th^ in RM 1 (Fig. 2A). This seems to indicate that negative contributions allow the RMs to better differentiate between sequences. One exception to this is Cys-containing peptides, which generally have lower response regardless of Cys being first or second in the motif (Fig. 3) but does not receive any extraordinary attention in any of the three RMs (Fig. 2). While the RMs do indeed provide new insight into the contribution of specific dimer motif occurrence, neither attention weights nor intensity data takes into account where in the peptide a particular dimer is located, and if the specific position matters when attempting to correlate peptide sequence and MS1 response.

### 2.3. Location, location, location?

To investigate if the specific position of dimer motifs affects the response, we selected the ten motifs with highest attention weights from RM 1-3 (Fig. S1-S3) as well as the ten motifs with highest (Fig. S4) and lowest (Fig. S5) mean MS1 response within the dataset and plotted the position-dependent mean MS1 response with the corresponding standard deviation. From the plots, eight representative motifs are shown below (Fig. 4). Common for all motifs at any position is the large variability of the MS1 intensities resulting in large standard deviations. As such, confidence intervals overlap, and no statistically significant differences were found. Not surprisingly, this illustrates that inclusion of any specific motif at a defined position within a peptide is insufficient for describing the complex phenomenon of overall variability in responses between equimolar peptides. While this data suggests no significant effect, there are, however, noticeable trends in the data that suggest the position of specific dimers does affect the MS1 response.

**Figure 4.**
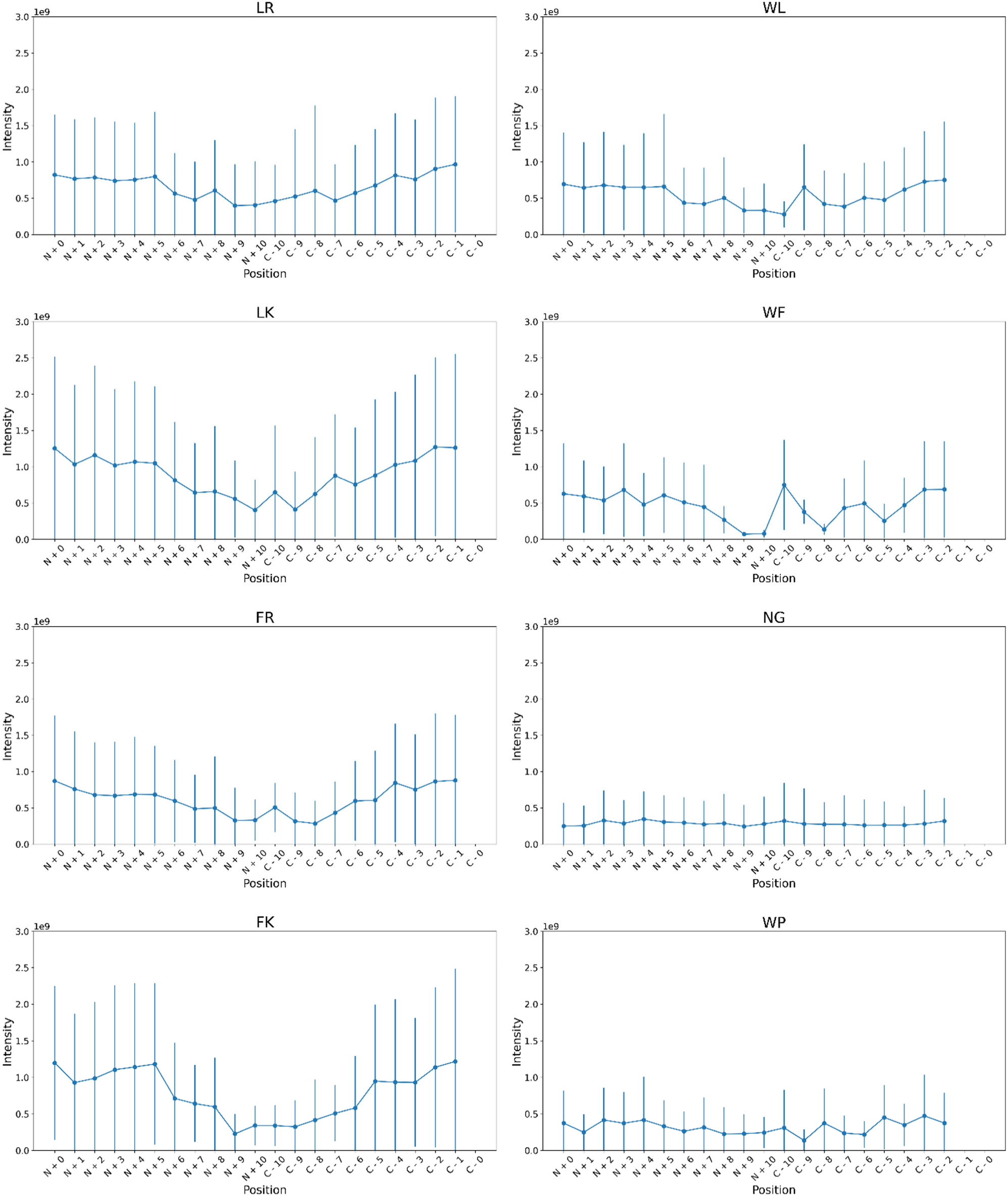
Mean intensity of eight representative dimer motifs (LR, LK, FR, FK, WL, WF, NG, WP) based on position (N-terminus to N + 10 AAs and C – 10 AAs to C-terminus for the first AA in the dimer) within the peptide sequence. Plots are shown as position-dependent mean intensity with standard deviations within the filtered dataset.

In general, dimers associated with high responding peptides and found important in RM 1 (e.g., LR, LK, FR, FK) seem to have higher mean responses when found in close vicinity (within five AAs) of peptide termini although high standard deviations lead to no statistically significant difference (Fig. 4, Fig. S1, Fig. S4). Moreover, dimers with Lys in the second position appear more dependent on position that dimers with Arg in the second position, indicating that not only does the inclusion of Lys appear to promote larger MS1 responses (Fig. 3), but having Lys adjacent to (particularly subsequent to) specific AAs towards the peptide termini is associated with higher responses. Interestingly, the data also suggests that there is negligible difference between peptide termini for this class of dimer motif. In contrast, dimer motifs associated with lower response but found of high relevance particularly in RM 2 and RM3 (e.g., WL, WF, NG, WP) are less affected by the position within the sequence (Fig. 4, Fig. S2, Fig. S3, Fig. S5). This observation is particularly evident for dimer motifs associated with the overall lowest responding peptides (Fig. S5), which includes Asn-Gly and a large number of Trp-, and Cys-, and His-containing peptides, indicating that any peptide including such motifs and/or AAs are likely to have lower responses than peptide which do not. This also correlates well with previous work investigating single AA occurrence, where particularly Trp, but also His and Cys, were identified as important to describe the relation between peptide sequence and MS1 response [27]. Nevertheless, combining AAs linked to low response (i.e., Trp, Cys, and His) with AAs associated with higher response (e.g., Leu, Ile, Lys, Phe) in a dimer motif (e.g., WL and WF) does, to some extent, alleviate the negative effect of the former on the MS1 response as well as increase positional dependency (Fig. 3, Fig. 4).

### 2.4. MS1 intensity prediction

The ability of the RMs to predict the MS1 response based on peptide sequence was investigated using the test subset of the filtered data. The performance was evaluated using the mean average percentage error (MAPE) for individual predictions and the overall agreement between predicted and real values using Pearson correlation coefficients (PCCs). The MAPE for the log-transformed MS1 intensity prediction were 11.5%, 12.2% and 10.8% with PCCs of 0.65, 0.64 and 0.63 for RM 1, 2, and 3, respectively (Table S1). This performance is comparable to but slightly poorer than performance metrics obtained in our previous work [27], where a similar model architecture considering only single amino acids obtained a MAPE of 10.5% and a PCC of 0.68). This could be explained by an increase in the number of variables, as our previous model considered peptide sequences as single AAs instead of all possible combination of two consecutive AAs, thereby increasing the combinatorial space from 21 to 401 variables. The higher complexity of the problem could have had a negative effect on the models’ performances. Indifferent to hyperparameters applied during training, the RMs all seem to be able to identify a relationship between sequences represented as dimers and the corresponding MS1 intensity values (Fig. 5). Moreover, the performances of the RMs may be impaired by the intrinsic variability within the dataset, owing to experimental factors and synthesizability aspects [13,27].

**Figure 5.**
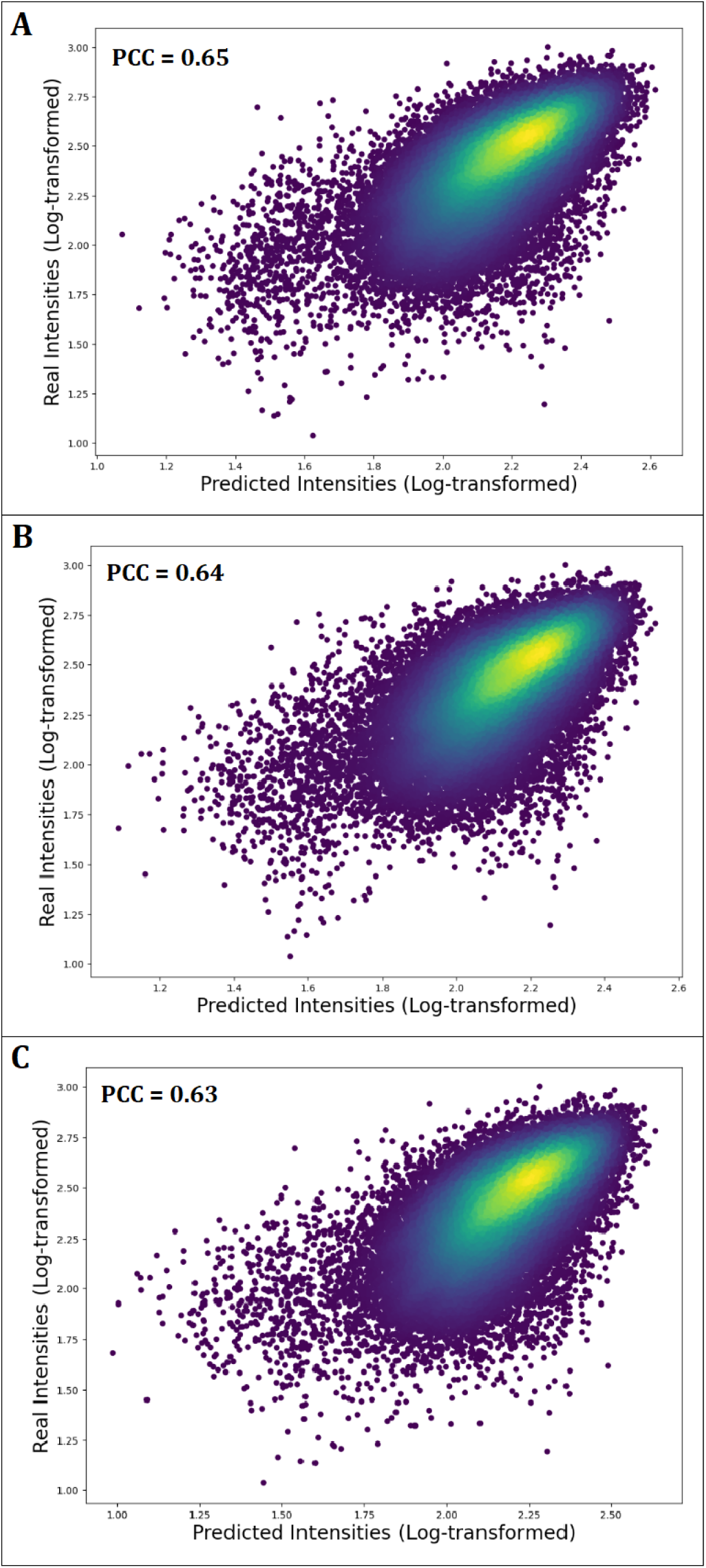
Scatter and density plot (from blue to yellow) of real versus predicted log-transformed MS1 intensities for A) Representative model 1; B) Representative model 2; and C) Representative model 3.

## 3. Conclusions

In this work, we present a deep learning model architecture for prediction of peptide MS1 intensity in LC-MS/MS analysis using a dimer motif peptide sequence representation by repurposing the largest equimolar dataset to date. The models presented in this work predicted MS1 intensity with a MAPE of ∼11% and a PCC of ∼0.64, indicating that the models are able to identify a correlation between peptide sequences and MS1 intensity output. Although the performance of the models was comparable, the dimer motifs identified as important by the built-in attention mechanism were considerably different. Three patterns in the identified motif were consistently observed. The first pattern represents the combination of bulky/aromatic hydrophobic AAs (Trp, Phe, Leu, Ile) followed by a cationic AA (Arg/Lys). The second pattern represents particularly aromatic AAs followed by bulky/aromatic hydrophobic AAs. The last pattern points particularly at Asn-Gly. Correlating identified motifs to mean intensities of peptides containing these motifs both globally and, in a position-dependent manner, further insight was obtained on how their inclusion affects the intensity output. Dimer motifs containing Trp as well as Asn-Gly were found to be associated with low responding peptides and indicated little or no positional bias. In contrast, motifs containing other bulky/aromatic hydrophobic AAs (i.e., Phe, Tyr, Ile, Leu) were found to be associated with high responding peptides; particularly when such an AA is followed by a similar AA or Lys. The impact of such motifs also appears to be more pronounced when found in the vicinity of peptide termini. Interestingly, the models identified motifs with Arg in the second position as more important than motifs with Lys in second position, but generally peptides with Lys in the second position have higher response. Together, these observations indicate that the models can more efficiently distinguish between peptides and predict their intensity by focusing on negative contributions. Ultimately, the findings of this study provide new insight on the understanding of peptide behavior during ESI-MS analysis. Moreover, the presented work also illustrates that DL is not only capable of identifying important contributors to MS1 response, but also has potential for developing methods that may accurately quantify peptides in complex mixtures without the need for tags, labels and custom standards. Nevertheless, more rigorous and comprehensive datasets as well as standardized workflows and data normalization strategies would be required to accomplish this.

## 4. Materials and methods

### 4.1. Datasets

The data used in this study was obtained from the PRIDE repository with the identifiers PXD004732[58], PXD010595[59], PXD021013[60]. In brief, the datasets were generated using equimolar pools of 1000 synthetic peptides each and originate from the development of Prosit [59,60] and ProteomeTools [58]. The dataset contains more than 4 million peptide identifications, representing over 1 million unique peptide sequences. The instrumentation used for the analyses were a Dionex 3000 HPLC system (Thermo Fisher Scientific) coupled online to an Orbitrap Fusion Lumos mass spectrometer (Thermo Fisher Scientific)[58–60]. The data was processed using MaxQuant [61] where specific, semi-specific, and unspecific enzyme setting were used, as well as, the proteases Trypsin, LysN and AspN. Repository data and metadata was extracted and processed using a custom Python (v.3.8.8) script, as previously described [27].

### 4.2. Data filtering and pre-processing

To reduce noise in the data and improve model performance, the data was initially filtered, as previously described [27]. In short, peptides were filtered based on quality and reproducibility criteria. The following exclusion criteria were defined:

- Posterior error probability (PEP) ≥ 0.01.
- Sequence identified in the reverse order (false positives).
- Listed as potential contaminants.
- Intensity value equal to zero.
- Identified as non-tryptic and not detected using a specific digestion setting.
- Duplicated peptides, keeping the median intensity MS output. Only identified once across all pools.
- Coefficient of variation (CV) ≥ 0.3 in replicate intensity measurements across pools.

To alleviate redundancy in the dataset, median peptide MS1 intensity across different pools was used as a representative response for individual peptides. The database was reduced to 179.222 unique specific/tryptic peptides after the filtering process. Specific/tryptic peptides refers to peptides that were identified during the processing of the MS raw data output (with MaxQuant) where a specific enzyme setting was used, and trypsin was the selected protease. Afterwards, the intensity values for each peptide sequence were log-transformed using the natural logarithm (Ln) and scaled between a range of values from 1 to 3. Lastly, the data was randomly split into training, validation, and test datasets. 90% of the data was used for the training dataset, from which 20% taken to build the validation dataset. The last 10% of the data was used as test dataset, to provide an unbiased evaluation of the final model.

### 4.3. Model Architecture

An encoder-decoder with attention mechanism was used in this study. The encoder was composed of one bidirectional recurrent neural network (BRNN) [62] with gated recurrent units (GRUs) [63]. The decoder had the same design as the encoder but with inclusion of a dense layer (Fig. 6). The BRNN with GRU for the encoder and decoder have the same number of units, while the dense layer has 1 unit corresponding to the predicted intensity values.

**Figure 6.**
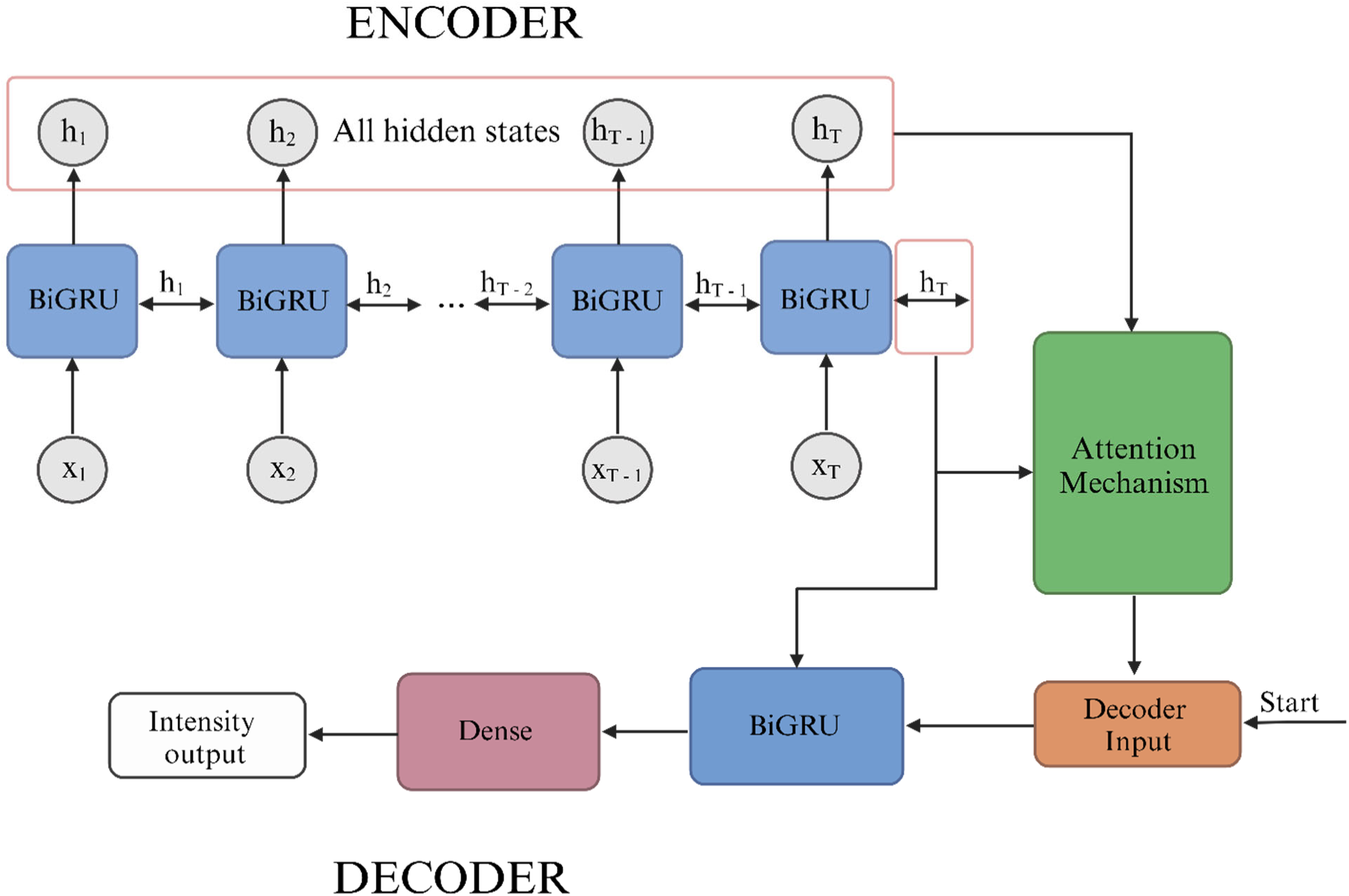
General scheme of the model architecture. The encoder consists of one BiGRU layer from which the attention mechanism takes all hidden states and the last hidden state to compute the context vector, which is then fed to the decoder together with the start character. The decoder consists of one BiGRU layer and a dense layer. The initial state of the decoder is the last hidden state of the encoder.

### 4.4. Training and testing

All the computational studies were done in Python (v.3.8.8) with TensorFlow[64] (v. 2.5.0) using the following libraries: Scikit-learn[65] (1.1.2), Statsmodels[66] (v. 0.13.2), Pandas[67] (v.1.4.4), Matplotlib[68] (v.3.5.2), Seaborn[69] (v.0.11.2), SciPy[70] (v.1.9.1), and NumPy[71] (v.1.23.1). The input data for the models were the one-hot encoded peptide sequences with a length ranging from 7 to 40 AAs, padding was applied to peptides with less than 40 AAs. AAs were grouped into k-mers of size 2, amounting a total of 401 possible AA combination including the padding character. The grouping of the AAs was done with a sliding window with a step of 1, meaning that all AAs were considered twice with the exception of the first and last AA in the peptide sequence. Thus, the dimension of the input data is batch size x 40 x 401. Different batch sizes, as well as number of units for the BRNN with GRU were used to generate different models presented in this work (Table S1).

To establish the relevancy of each dimer motif, the attention weights of the models were determined for each of the 401 possible combinations including padding. The attention weights are averaged for the repeated dimers in the same sequence (if applicable) and then across all peptide sequences evaluated, generating an overall mean on the attention weight for each dimer motif. The values of those attention weights will rank the dimer motifs from the highest to the lowest relevancy within to each model. The predicted MS1 intensity output was used to evaluate the models’ performance as previously described [27]. The loss function used to train the models was the mean squared error (MSE) [72,73]. while the mean absolute error (MAE) was used as accuracy measurement. The final results of the evaluation of the models were expressed using the mean absolute percentage error (MAPE) [74,75]. Adam [76] was the optimizer selected during the training process of the models. The models were trained on NVIDIA Quadro T2000 GPU for 5 to 10 epochs.

## 5. Data availability

The data used in this study was obtained from the PRIDE repository with the identifiers PXD004732, PXD010595, and PXD021013.

## 6. Author contributions

**Naim Abdul-Khalek:** Conceptualization, Methodology, Software, Validation, Formal analysis, Investigation, Data Curation, Writing - Original Draft, Writing - Review & Editing, Visualization, Project administration. **Reinhard Wimmer:** Conceptualization, Resources, Writing - Review & Editing, Supervision. **Michael Toft Overgaard:** Conceptualization, Resources, Writing - Review & Editing, Supervision, Funding acquisition. **Simon Gregersen Echers:** Conceptualization, Methodology, Resources, Writing - Original Draft, Writing - Review & Editing, Supervision, Project administration, Funding acquisition.

