## Supplementary information for "Decoding the Impact of Neighboring Amino Acid on ESI-MS Intensity Output through Deep Learning"

### Supplementary information
### for

#### Decoding the Impact of Neighboring Amino Acid on ESI-MS Intensity Output through Deep Learning

Naim Abdul-Khalek[1], Reinhard Wimmer[1], Michael Toft Overgaard[1], Simon Gregersen Echers[1, *]

[1] Department of Chemistry and Bioscience, Aalborg University, Fredrik Bajers Vej 7H, Aalborg 9220, Denmark.

 (Gregersen Echers S)

**Running title:** *Abdul-Khalek, N et al. / Dimers and peptide MS response by Deep Learning*

**Table S1**. Performance metrics and hyperparameters for the three representative models, showing performance by mean average percentage error (MAPE, %), Pearson correlation coefficient (PCC) as well as batch size, units, epochs, and output range applied during training for the individual representative models.

| | MAPE (%) Log-Transformed | PCC | Batch Size | Units | Epochs | Output range of values |
|---|---|---|---|---|---|---|
| **Representative Model 1** | 11.2 | 0.65 | 128 | 32 | 10 | 1-3 |
| **Representative Model 2** | 12.2 | 0.64 | 512 | 256 | 5 | 1-3 |
| **Representative Model 3** | 10.8 | 0.63 | 64 | 32 | 5 | 1-3 |

**Table S2.** Occurences of AA dimer motifs within the dataset. In the table, all occurences are included across all peptides and a specifc motif may be present multiple times within the same peptide sequence.

| | | First Amino Acid | | | | | | | | | | | | | | | | | | | |
|---|---|---|---|---|---|---|---|---|---|---|---|---|---|---|---|---|---|---|---|---|---|
| | | **A** | **C** | **D** | **E** | **F** | **G** | **H** | **I** | **K** | **L** | **M** | **N** | **P** | **Q** | **R** | **S** | **T** | **V** | **W** | **Y** |
| **Second Amino Acid** | **A** | 18444 | 2984 | 7525 | 14748 | 4188 | 12247 | 2886 | 5516 | 3631 | 16878 | 2974 | 5057 | 14999 | 9643 | 3644 | 13386 | 9080 | 10204 | 1699 | 3187 |
| | **C** | 3420 | 1461 | 2655 | 5273 | 2033 | 3269 | 1464 | 2669 | 1149 | 5229 | 677 | 1913 | 3420 | 2714 | 1003 | 4641 | 3004 | 3415 | 483 | 1554 |
| | **D** | 7520 | 2514 | 5606 | 11028 | 3808 | 7705 | 1809 | 4789 | 3166 | 12162 | 2311 | 3689 | 7868 | 6585 | 2519 | 9323 | 5902 | 7075 | 1384 | 3057 |
| | **E** | 13502 | 3439 | 8390 | 20962 | 5058 | 12319 | 3012 | 6748 | 5104 | 20732 | 3773 | 6021 | 13502 | 11654 | 3712 | 13648 | 9311 | 10009 | 1852 | 4914 |
| | **F** | 7426 | 1674 | 5213 | 5404 | 2895 | 5623 | 2401 | 3768 | 1447 | 6862 | 1134 | 3703 | 5211 | 3307 | 1608 | 7607 | 4998 | 4982 | 592 | 2562 |
| | **G** | 13097 | 5537 | 7504 | 11487 | 4996 | 12933 | 3206 | 4909 | 2771 | 15871 | 2271 | 3458 | 15713 | 8156 | 3430 | 14075 | 9587 | 8458 | 1673 | 3936 |
| | **H** | 3360 | 1210 | 2400 | 3551 | 2244 | 3451 | 1461 | 3763 | 1012 | 6030 | 673 | 2044 | 3371 | 3108 | 1158 | 4892 | 3288 | 3517 | 506 | 1507 |
| | **I** | 6137 | 1907 | 6467 | 8431 | 3815 | 6191 | 2447 | 5092 | 2358 | 8875 | 1386 | 5085 | 5022 | 5164 | 2121 | 7450 | 5263 | 6185 | 905 | 3033 |
| | **K** | 7790 | 3046 | 6148 | 14192 | 4735 | 10456 | 2764 | 6567 | 2690 | 15378 | 2722 | 5215 | 7929 | 7986 | 2158 | 10395 | 6311 | 8252 | 1860 | 4152 |
| | **L** | 18484 | 5005 | 13525 | 17926 | 8805 | 15451 | 6946 | 10207 | 4547 | 24819 | 3668 | 10053 | 15419 | 13329 | 5040 | 21853 | 13203 | 14307 | 2108 | 6415 |
| | **M** | 2465 | 625 | 1954 | 3427 | 1130 | 2048 | 816 | 1580 | 858 | 3140 | 723 | 1569 | 1908 | 1987 | 725 | 2591 | 1837 | 2116 | 439 | 891 |
| | **N** | 4552 | 2045 | 4075 | 8567 | 3375 | 5419 | 1646 | 4376 | 1986 | 8703 | 1708 | 3670 | 4574 | 4737 | 1760 | 6593 | 4001 | 5199 | 1009 | 2662 |
| | **P** | 13075 | 4134 | 7966 | 9839 | 5476 | 13115 | 3737 | 6571 | 6606 | 17525 | 2196 | 6050 | 16191 | 7800 | 6718 | 18716 | 10264 | 9824 | 1287 | 3493 |
| | **Q** | 8884 | 2633 | 4984 | 9000 | 5043 | 7427 | 3829 | 6351 | 2196 | 16918 | 1837 | 4672 | 8740 | 8077 | 2086 | 11055 | 6424 | 7323 | 1226 | 3542 |
| | **R** | 10568 | 3466 | 5861 | 10784 | 5726 | 10617 | 3656 | 6382 | 2228 | 17428 | 2016 | 4577 | 12386 | 9093 | 2906 | 12756 | 6858 | 8331 | 2013 | 4347 |
| | **S** | 14826 | 4921 | 10156 | 11423 | 8914 | 14280 | 4970 | 9018 | 3516 | 21194 | 2797 | 7849 | 16810 | 8319 | 3610 | 22936 | 10843 | 12028 | 1752 | 5350 |
| | **T** | 8652 | 2477 | 6155 | 8883 | 4780 | 8469 | 4104 | 5736 | 2581 | 12951 | 2080 | 4806 | 8690 | 6054 | 2420 | 10633 | 6764 | 8343 | 1349 | 3283 |
| | **V** | 11124 | 2722 | 7833 | 10963 | 4249 | 8432 | 2960 | 5218 | 3032 | 12328 | 2310 | 5279 | 10546 | 7660 | 2749 | 11339 | 8323 | 8538 | 1276 | 3538 |
| | **W** | 1840 | 573 | 1260 | 1553 | 810 | 1665 | 644 | 870 | 459 | 2041 | 303 | 922 | 1902 | 1177 | 576 | 2319 | 1326 | 1269 | 306 | 640 |
| | **Y** | 3608 | 1294 | 3608 | 4113 | 2600 | 3929 | 1641 | 3144 | 1395 | 5783 | 884 | 2859 | 4974 | 2723 | 1231 | 5163 | 3314 | 3635 | 617 | 2154 |

**Table S3**. Attention weights of AA dimer motifs for representative model 1.

| | | First Amino Acid | | | | | | | | | | | | | | | | | | | |
|---|---|---|---|---|---|---|---|---|---|---|---|---|---|---|---|---|---|---|---|---|---|
| | | **A** | **C** | **D** | **E** | **F** | **G** | **H** | **I** | **K** | **L** | **M** | **N** | **P** | **Q** | **R** | **S** | **T** | **V** | **W** | **Y** |
| **Second Amino Acid** | **A** | 0,0502 | 0,0551 | 0,0466 | 0,0458 | 0,0671 | 0,0463 | 0,0526 | 0,0747 | 0,0484 | 0,0718 | 0,0518 | 0,0461 | 0,0614 | 0,0508 | 0,0480 | 0,0547 | 0,0444 | 0,0590 | 0,1019 | 0,0569 |
| | **C** | 0,0556 | 0,0847 | 0,0555 | 0,0457 | 0,0858 | 0,0553 | 0,0605 | 0,0940 | 0,0532 | 0,0828 | 0,0592 | 0,0589 | 0,0669 | 0,0570 | 0,0380 | 0,0524 | 0,0623 | 0,0740 | 0,1052 | 0,0588 |
| | **D** | 0,0503 | 0,0570 | 0,0575 | 0,0491 | 0,0734 | 0,0431 | 0,0503 | 0,0760 | 0,0426 | 0,0784 | 0,0466 | 0,0522 | 0,0577 | 0,0592 | 0,0385 | 0,0428 | 0,0454 | 0,0602 | 0,0845 | 0,0691 |
| | **E** | 0,0491 | 0,0502 | 0,0616 | 0,0431 | 0,0812 | 0,0431 | 0,0384 | 0,0686 | 0,0404 | 0,0760 | 0,0604 | 0,0466 | 0,0735 | 0,0385 | 0,0408 | 0,0545 | 0,0480 | 0,0572 | 0,0961 | 0,0853 |
| | **F** | 0,0438 | 0,0511 | 0,0492 | 0,0513 | 0,0748 | 0,0462 | 0,0490 | 0,0737 | 0,0497 | 0,0728 | 0,0680 | 0,0531 | 0,0625 | 0,0448 | 0,0556 | 0,0515 | 0,0522 | 0,0647 | 0,0979 | 0,0599 |
| | **G** | 0,0477 | 0,0584 | 0,0463 | 0,0534 | 0,0833 | 0,0414 | 0,0337 | 0,0709 | 0,0562 | 0,0788 | 0,0567 | 0,1699 | 0,0540 | 0,0417 | 0,0417 | 0,0412 | 0,0388 | 0,0644 | 0,0729 | 0,0632 |
| | **H** | 0,0409 | 0,0427 | 0,0572 | 0,0571 | 0,0761 | 0,0655 | 0,0734 | 0,0824 | 0,0856 | 0,0693 | 0,0699 | 0,0876 | 0,0764 | 0,0586 | 0,0578 | 0,0496 | 0,0527 | 0,0709 | 0,0929 | 0,0690 |
| | **I** | 0,0595 | 0,0514 | 0,0595 | 0,0510 | 0,0866 | 0,0528 | 0,0491 | 0,0797 | 0,0547 | 0,0730 | 0,0483 | 0,0613 | 0,0713 | 0,0467 | 0,0539 | 0,0521 | 0,0554 | 0,0567 | 0,0883 | 0,0828 |
| | **K** | 0,0760 | 0,0948 | 0,0711 | 0,0695 | 0,1616 | 0,0771 | 0,0751 | 0,1290 | 0,0961 | 0,1496 | 0,0907 | 0,0656 | 0,1024 | 0,0933 | 0,0773 | 0,0832 | 0,0829 | 0,1137 | 0,1558 | 0,1385 |
| | **L** | 0,0499 | 0,0573 | 0,0481 | 0,0549 | 0,0697 | 0,0534 | 0,0528 | 0,0690 | 0,0578 | 0,0664 | 0,0536 | 0,0533 | 0,0531 | 0,0522 | 0,0486 | 0,0496 | 0,0494 | 0,0659 | 0,0907 | 0,0619 |
| | **M** | 0,0645 | 0,0503 | 0,0575 | 0,0545 | 0,0923 | 0,0426 | 0,0665 | 0,0681 | 0,0481 | 0,0659 | 0,0483 | 0,0650 | 0,0785 | 0,0510 | 0,0601 | 0,0502 | 0,0517 | 0,0552 | 0,0810 | 0,0623 |
| | **N** | 0,0528 | 0,0545 | 0,0574 | 0,0417 | 0,0742 | 0,0425 | 0,0578 | 0,0881 | 0,0468 | 0,0811 | 0,0559 | 0,0553 | 0,0744 | 0,0463 | 0,0489 | 0,0471 | 0,0490 | 0,0629 | 0,0885 | 0,0666 |
| | **P** | 0,0422 | 0,0688 | 0,0506 | 0,0540 | 0,0819 | 0,0429 | 0,0493 | 0,0672 | 0,0308 | 0,0706 | 0,0425 | 0,0554 | 0,0526 | 0,0464 | 0,0425 | 0,0376 | 0,0466 | 0,0542 | 0,1061 | 0,0620 |
| | **Q** | 0,0453 | 0,0487 | 0,0568 | 0,0508 | 0,0821 | 0,0481 | 0,0471 | 0,0697 | 0,0445 | 0,0772 | 0,0491 | 0,0485 | 0,0695 | 0,0473 | 0,0441 | 0,0443 | 0,0486 | 0,0614 | 0,0977 | 0,0591 |
| | **R** | 0,1224 | 0,0947 | 0,1100 | 0,1388 | 0,1944 | 0,1020 | 0,1076 | 0,1851 | 0,1513 | 0,1865 | 0,1204 | 0,1087 | 0,1421 | 0,1153 | 0,1234 | 0,1171 | 0,1155 | 0,1630 | 0,1872 | 0,1596 |
| | **S** | 0,0481 | 0,0743 | 0,0462 | 0,0437 | 0,0780 | 0,0433 | 0,0548 | 0,0697 | 0,0331 | 0,0749 | 0,0502 | 0,0496 | 0,0688 | 0,0540 | 0,0401 | 0,0424 | 0,0463 | 0,0642 | 0,0854 | 0,0657 |
| | **T** | 0,0471 | 0,0612 | 0,0546 | 0,0456 | 0,0752 | 0,0473 | 0,0392 | 0,0738 | 0,0557 | 0,0784 | 0,0582 | 0,0481 | 0,0685 | 0,0541 | 0,0510 | 0,0437 | 0,0444 | 0,0631 | 0,0857 | 0,0634 |
| | **V** | 0,0591 | 0,0525 | 0,0541 | 0,0530 | 0,0758 | 0,0412 | 0,0460 | 0,0624 | 0,0494 | 0,0768 | 0,0674 | 0,0583 | 0,0600 | 0,0507 | 0,0631 | 0,0563 | 0,0598 | 0,0628 | 0,0802 | 0,0553 |
| | **W** | 0,0635 | 0,0483 | 0,0599 | 0,0641 | 0,0934 | 0,0553 | 0,0661 | 0,0840 | 0,0658 | 0,0886 | 0,0491 | 0,0614 | 0,0756 | 0,0644 | 0,0565 | 0,0634 | 0,0512 | 0,0901 | 0,0716 | 0,0900 |
| | **Y** | 0,0606 | 0,0730 | 0,0548 | 0,0545 | 0,0704 | 0,0457 | 0,0411 | 0,0718 | 0,0488 | 0,0640 | 0,0585 | 0,0545 | 0,0505 | 0,0511 | 0,0447 | 0,0519 | 0,0500 | 0,0609 | 0,0904 | 0,0716 |

**Table S4**. Attention weights of AA dimer motifs for representative model 2.

| | | First Amino Acid | | | | | | | | | | | | | | | | | | | |
| --- | --- | --- | --- | --- | --- | --- | --- | --- | --- | --- | --- | --- | --- | --- | --- | --- | --- | --- | --- | --- | --- |
| | | **A** | **C** | **D** | **E** | **F** | **G** | **H** | **I** | **K** | **L** | **M** | **N** | **P** | **Q** | **R** | **S** | **T** | **V** | **W** | **Y** |
| **Second Amino Acid** | **A** | 0,0549 | 0,0597 | 0,0629 | 0,0675 | 0,1305 | 0,0544 | 0,0526 | 0,0997 | 0,0587 | 0,0822 | 0,0753 | 0,0648 | 0,0559 | 0,0576 | 0,0373 | 0,0514 | 0,0440 | 0,0653 | 0,1682 | 0,1164 |
| | **C** | 0,0342 | 0,0522 | 0,0426 | 0,0371 | 0,1004 | 0,0543 | 0,0332 | 0,0721 | 0,0374 | 0,0839 | 0,0498 | 0,0621 | 0,0569 | 0,0481 | 0,0322 | 0,0440 | 0,0508 | 0,0652 | 0,0916 | 0,0940 |
| | **D** | 0,0448 | 0,0645 | 0,0585 | 0,0454 | 0,1133 | 0,0504 | 0,0427 | 0,0896 | 0,0381 | 0,0963 | 0,0713 | 0,0495 | 0,0690 | 0,0454 | 0,0404 | 0,0482 | 0,0541 | 0,0658 | 0,1521 | 0,0898 |
| | **E** | 0,0477 | 0,0682 | 0,0618 | 0,0558 | 0,0884 | 0,0546 | 0,0477 | 0,0883 | 0,0347 | 0,0916 | 0,0886 | 0,0612 | 0,0615 | 0,0559 | 0,0370 | 0,0431 | 0,0524 | 0,0828 | 0,1702 | 0,0962 |
| | **F** | 0,0859 | 0,1115 | 0,0925 | 0,0931 | 0,1616 | 0,0813 | 0,0797 | 0,1114 | 0,0553 | 0,1234 | 0,1302 | 0,1039 | 0,1170 | 0,0920 | 0,0664 | 0,0931 | 0,0849 | 0,0993 | 0,1851 | 0,1136 |
| | **G** | 0,0480 | 0,0570 | 0,0530 | 0,0557 | 0,1004 | 0,0403 | 0,0478 | 0,0788 | 0,0369 | 0,0808 | 0,0573 | 0,1198 | 0,0626 | 0,0458 | 0,0417 | 0,0367 | 0,0382 | 0,0534 | 0,1207 | 0,1015 |
| | **H** | 0,0294 | 0,0461 | 0,0380 | 0,0472 | 0,0752 | 0,0386 | 0,0556 | 0,0651 | 0,0474 | 0,0647 | 0,0488 | 0,0476 | 0,0469 | 0,0539 | 0,0333 | 0,0353 | 0,0384 | 0,0470 | 0,1180 | 0,0595 |
| | **I** | 0,0819 | 0,1140 | 0,1188 | 0,0971 | 0,1383 | 0,0795 | 0,0813 | 0,0922 | 0,0689 | 0,1096 | 0,0985 | 0,1047 | 0,0861 | 0,0909 | 0,0687 | 0,0940 | 0,0938 | 0,0874 | 0,1830 | 0,1195 |
| | **K** | 0,0176 | 0,0245 | 0,0305 | 0,0240 | 0,0658 | 0,0203 | 0,0235 | 0,0424 | 0,0179 | 0,0423 | 0,0378 | 0,0194 | 0,0242 | 0,0227 | 0,0180 | 0,0192 | 0,0210 | 0,0306 | 0,0754 | 0,0318 |
| | **L** | 0,0830 | 0,1022 | 0,1245 | 0,0878 | 0,1460 | 0,0945 | 0,0674 | 0,1211 | 0,0790 | 0,1151 | 0,0998 | 0,1092 | 0,0931 | 0,0867 | 0,0844 | 0,0856 | 0,0798 | 0,0928 | 0,1886 | 0,1364 |
| | **M** | 0,0605 | 0,0657 | 0,0771 | 0,0695 | 0,1348 | 0,0741 | 0,0495 | 0,0775 | 0,0533 | 0,0964 | 0,0741 | 0,1024 | 0,0758 | 0,0705 | 0,0483 | 0,0624 | 0,0608 | 0,0762 | 0,1258 | 0,0930 |
| | **N** | 0,0511 | 0,0457 | 0,0649 | 0,0430 | 0,0991 | 0,0453 | 0,0473 | 0,0786 | 0,0331 | 0,0791 | 0,0695 | 0,0440 | 0,0654 | 0,0401 | 0,0418 | 0,0446 | 0,0529 | 0,0668 | 0,1177 | 0,0942 |
| | **P** | 0,0654 | 0,0737 | 0,0877 | 0,0870 | 0,1333 | 0,0624 | 0,0617 | 0,1082 | 0,0605 | 0,1098 | 0,1143 | 0,0943 | 0,0702 | 0,0881 | 0,0793 | 0,0731 | 0,0809 | 0,0942 | 0,1787 | 0,1326 |
| | **Q** | 0,0423 | 0,0520 | 0,0578 | 0,0477 | 0,0802 | 0,0432 | 0,0454 | 0,1116 | 0,0328 | 0,0848 | 0,0793 | 0,0586 | 0,0635 | 0,0493 | 0,0384 | 0,0449 | 0,0432 | 0,0637 | 0,1186 | 0,0839 |
| | **R** | 0,0201 | 0,0268 | 0,0261 | 0,0226 | 0,0721 | 0,0260 | 0,0335 | 0,0445 | 0,0310 | 0,0441 | 0,0390 | 0,0257 | 0,0303 | 0,0180 | 0,0218 | 0,0203 | 0,0259 | 0,0377 | 0,0828 | 0,0499 |
| | **S** | 0,0399 | 0,0460 | 0,0462 | 0,0415 | 0,1020 | 0,0391 | 0,0354 | 0,0880 | 0,0316 | 0,0854 | 0,0531 | 0,0469 | 0,0693 | 0,0427 | 0,0375 | 0,0391 | 0,0461 | 0,0551 | 0,1076 | 0,0998 |
| | **T** | 0,0494 | 0,0677 | 0,0588 | 0,0491 | 0,1130 | 0,0479 | 0,0330 | 0,1055 | 0,0359 | 0,0858 | 0,0805 | 0,0569 | 0,0678 | 0,0463 | 0,0549 | 0,0412 | 0,0543 | 0,0622 | 0,1526 | 0,0661 |
| | **V** | 0,0759 | 0,0800 | 0,0897 | 0,0759 | 0,1160 | 0,0760 | 0,0499 | 0,1013 | 0,0528 | 0,1025 | 0,0882 | 0,0776 | 0,0747 | 0,0651 | 0,0602 | 0,0717 | 0,0736 | 0,1017 | 0,1727 | 0,1319 |
| | **W** | 0,1241 | 0,1567 | 0,1473 | 0,0905 | 0,1347 | 0,1177 | 0,0627 | 0,1110 | 0,0967 | 0,1251 | 0,1163 | 0,1080 | 0,1183 | 0,1192 | 0,0948 | 0,0994 | 0,1025 | 0,1227 | 0,1546 | 0,1379 |
| | **Y** | 0,0543 | 0,0948 | 0,0953 | 0,0721 | 0,1217 | 0,0680 | 0,0375 | 0,0897 | 0,0628 | 0,1114 | 0,0955 | 0,0895 | 0,0752 | 0,0699 | 0,0445 | 0,0810 | 0,0685 | 0,0787 | 0,1263 | 0,1260 |

**Table S5**. Attention weights of AA dimer motifs for representative model 3.

| | | First Amino Acid | | | | | | | | | | | | | | | | | | | |
|---|---|---|---|---|---|---|---|---|---|---|---|---|---|---|---|---|---|---|---|---|---|
| | | **A** | **C** | **D** | **E** | **F** | **G** | **H** | **I** | **K** | **L** | **M** | **N** | **P** | **Q** | **R** | **S** | **T** | **V** | **W** | **Y** |
| **Second Amino Acid** | **A** | 0,0565 | 0,0660 | 0,0484 | 0,0445 | 0,0970 | 0,0426 | 0,0613 | 0,0832 | 0,0519 | 0,0900 | 0,0628 | 0,0541 | 0,0646 | 0,0462 | 0,0422 | 0,0550 | 0,0428 | 0,0554 | 0,1500 | 0,0630 |
| | **C** | 0,0574 | 0,1241 | 0,0618 | 0,0474 | 0,1142 | 0,0658 | 0,0625 | 0,0797 | 0,0431 | 0,0959 | 0,0686 | 0,0909 | 0,0902 | 0,0588 | 0,0498 | 0,0708 | 0,0780 | 0,0751 | 0,1142 | 0,0678 |
| | **D** | 0,0435 | 0,0631 | 0,0520 | 0,0503 | 0,0782 | 0,0432 | 0,0566 | 0,0790 | 0,0413 | 0,0890 | 0,0645 | 0,0528 | 0,0644 | 0,0484 | 0,0522 | 0,0429 | 0,0456 | 0,0670 | 0,1126 | 0,0793 |
| | **E** | 0,0414 | 0,0632 | 0,0593 | 0,0426 | 0,0866 | 0,0380 | 0,0400 | 0,0863 | 0,0288 | 0,0945 | 0,0575 | 0,0468 | 0,0843 | 0,0462 | 0,0361 | 0,0483 | 0,0433 | 0,0605 | 0,1392 | 0,1006 |
| | **F** | 0,0463 | 0,1071 | 0,0611 | 0,0560 | 0,1054 | 0,0539 | 0,0711 | 0,0901 | 0,0531 | 0,0946 | 0,0964 | 0,0687 | 0,0843 | 0,0541 | 0,0639 | 0,0662 | 0,0711 | 0,0830 | 0,1581 | 0,0772 |
| | **G** | 0,0462 | 0,0911 | 0,0533 | 0,0423 | 0,1036 | 0,0376 | 0,0286 | 0,0786 | 0,0467 | 0,0894 | 0,0693 | 0,2486 | 0,0642 | 0,0389 | 0,0414 | 0,0320 | 0,0332 | 0,0629 | 0,1105 | 0,0607 |
| | **H** | 0,0373 | 0,0508 | 0,0635 | 0,0468 | 0,0780 | 0,0625 | 0,1106 | 0,0874 | 0,0881 | 0,0753 | 0,0712 | 0,0974 | 0,0856 | 0,0687 | 0,0794 | 0,0498 | 0,0606 | 0,0725 | 0,1081 | 0,0667 |
| | **I** | 0,0726 | 0,1089 | 0,0652 | 0,0576 | 0,1103 | 0,0609 | 0,0800 | 0,1030 | 0,0683 | 0,0926 | 0,0745 | 0,0713 | 0,0643 | 0,0581 | 0,0625 | 0,0532 | 0,0600 | 0,0693 | 0,1348 | 0,1087 |
| | **K** | 0,0455 | 0,0516 | 0,0368 | 0,0373 | 0,1375 | 0,0443 | 0,0632 | 0,0989 | 0,0792 | 0,1093 | 0,0583 | 0,0352 | 0,0782 | 0,0623 | 0,0756 | 0,0460 | 0,0428 | 0,0682 | 0,1337 | 0,0923 |
| | **L** | 0,0600 | 0,0848 | 0,0614 | 0,0560 | 0,0973 | 0,0582 | 0,0872 | 0,0872 | 0,0598 | 0,0982 | 0,0717 | 0,0592 | 0,0640 | 0,0573 | 0,0693 | 0,0590 | 0,0595 | 0,0670 | 0,1335 | 0,0813 |
| | **M** | 0,0702 | 0,0676 | 0,0738 | 0,0629 | 0,1118 | 0,0469 | 0,0644 | 0,0785 | 0,0501 | 0,0815 | 0,0614 | 0,0735 | 0,0910 | 0,0616 | 0,0675 | 0,0607 | 0,0563 | 0,0654 | 0,1073 | 0,0764 |
| | **N** | 0,0476 | 0,0793 | 0,0631 | 0,0358 | 0,0878 | 0,0427 | 0,0707 | 0,0873 | 0,0355 | 0,0886 | 0,0671 | 0,0516 | 0,0784 | 0,0419 | 0,0500 | 0,0382 | 0,0456 | 0,0562 | 0,1183 | 0,0651 |
| | **P** | 0,0456 | 0,1011 | 0,0624 | 0,0483 | 0,1181 | 0,0503 | 0,0742 | 0,0873 | 0,0432 | 0,0809 | 0,0599 | 0,0622 | 0,0757 | 0,0467 | 0,0688 | 0,0372 | 0,0571 | 0,0652 | 0,1543 | 0,0881 |
| | **Q** | 0,0348 | 0,0666 | 0,0494 | 0,0482 | 0,1005 | 0,0478 | 0,0764 | 0,0758 | 0,0445 | 0,0879 | 0,0599 | 0,0444 | 0,0691 | 0,0442 | 0,0481 | 0,0426 | 0,0451 | 0,0625 | 0,1216 | 0,0514 |
| | **R** | 0,0806 | 0,0717 | 0,0743 | 0,0821 | 0,1932 | 0,0859 | 0,1171 | 0,1641 | 0,1337 | 0,1765 | 0,0935 | 0,0838 | 0,1140 | 0,0774 | 0,1365 | 0,0867 | 0,0923 | 0,1216 | 0,1857 | 0,1215 |
| | **S** | 0,0471 | 0,0883 | 0,0423 | 0,0480 | 0,1033 | 0,0407 | 0,0672 | 0,0778 | 0,0294 | 0,0873 | 0,0520 | 0,0459 | 0,0786 | 0,0468 | 0,0431 | 0,0381 | 0,0441 | 0,0722 | 0,1238 | 0,0920 |
| | **T** | 0,0463 | 0,0886 | 0,0532 | 0,0507 | 0,1010 | 0,0475 | 0,0561 | 0,0848 | 0,0590 | 0,0960 | 0,0708 | 0,0472 | 0,0694 | 0,0533 | 0,0551 | 0,0451 | 0,0412 | 0,0647 | 0,0947 | 0,0721 |
| | **V** | 0,0774 | 0,0747 | 0,0653 | 0,0589 | 0,0993 | 0,0480 | 0,0693 | 0,0786 | 0,0583 | 0,0988 | 0,0829 | 0,0619 | 0,0801 | 0,0593 | 0,0754 | 0,0610 | 0,0673 | 0,0804 | 0,1150 | 0,0838 |
| | **W** | 0,0867 | 0,1149 | 0,0998 | 0,0835 | 0,1074 | 0,0682 | 0,0774 | 0,1097 | 0,0893 | 0,1186 | 0,0720 | 0,0845 | 0,1015 | 0,0849 | 0,0860 | 0,0727 | 0,0845 | 0,0937 | 0,1197 | 0,1019 |
| | **Y** | 0,0518 | 0,0862 | 0,0669 | 0,0520 | 0,0980 | 0,0554 | 0,0475 | 0,0881 | 0,0578 | 0,0823 | 0,0690 | 0,0648 | 0,0781 | 0,0624 | 0,0591 | 0,0554 | 0,0699 | 0,0760 | 0,1089 | 0,0967 |

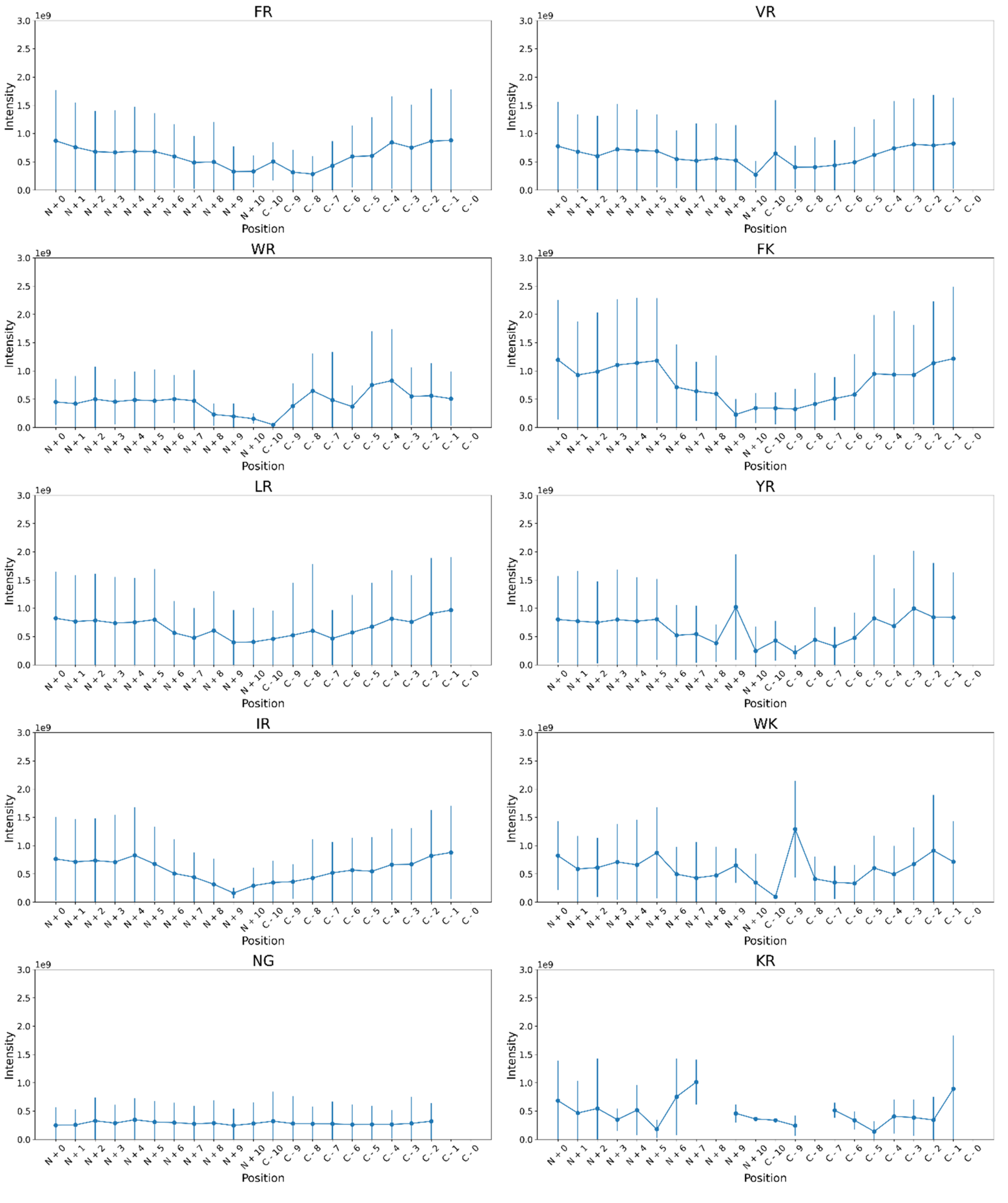


Figure S1: Mean intensity of top 10 dimer motifs from RM 1 based on position (N-terminus to N + 10 AAs and C – 10 AAs to C-terminus for the first AA in the dimer) within the peptide sequence. Plots are shown as position-dependent mean intensity with standard deviations within the filtered dataset.

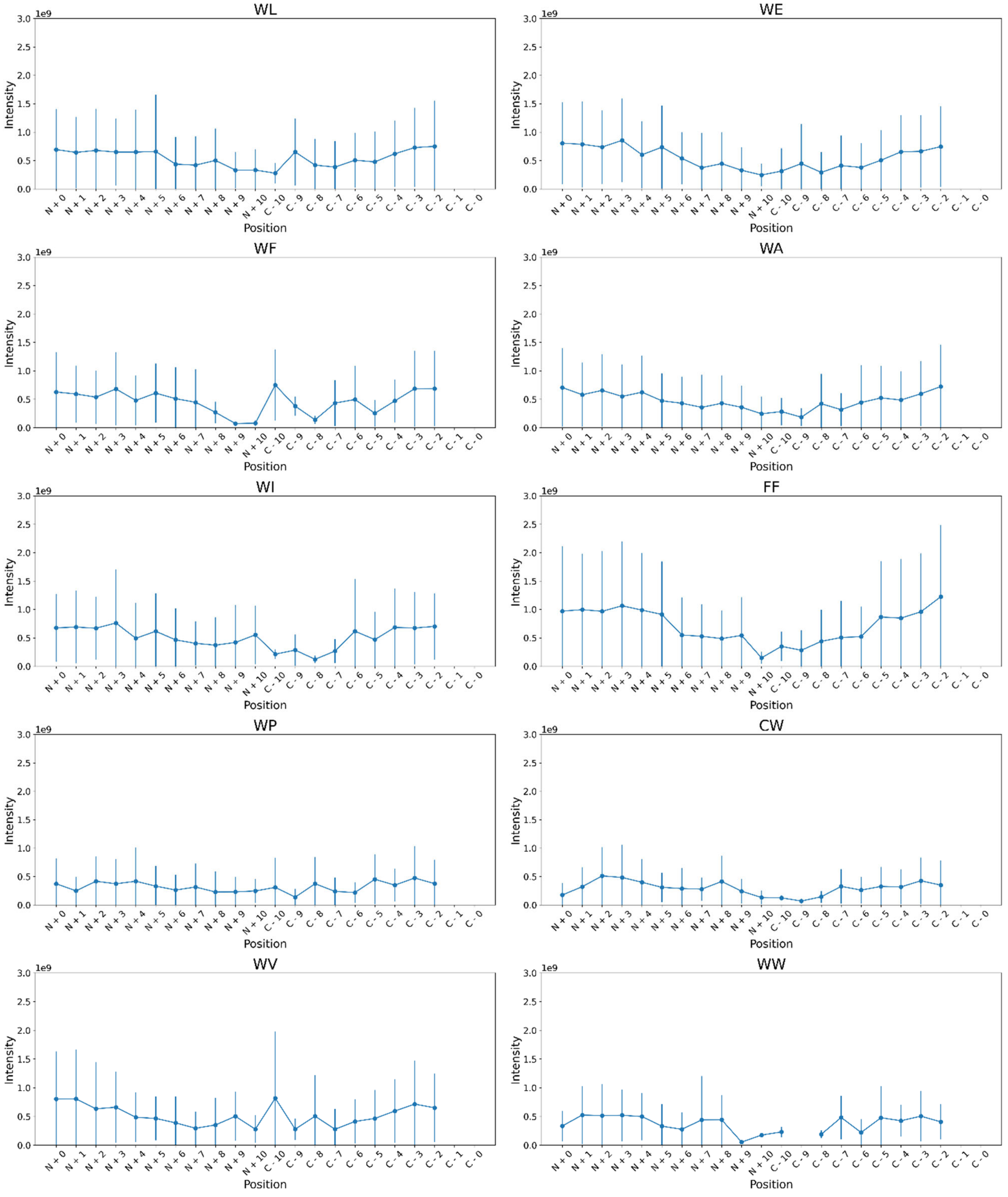


Figure S2: Mean intensity of top 10 dimer motifs from RM 2 based on position (N-terminus to N + 10 AAs and C – 10 AAs to C-terminus for the first AA in the dimer) within the peptide sequence. Plots are shown as position-dependent mean intensity with standard deviations within the filtered dataset.

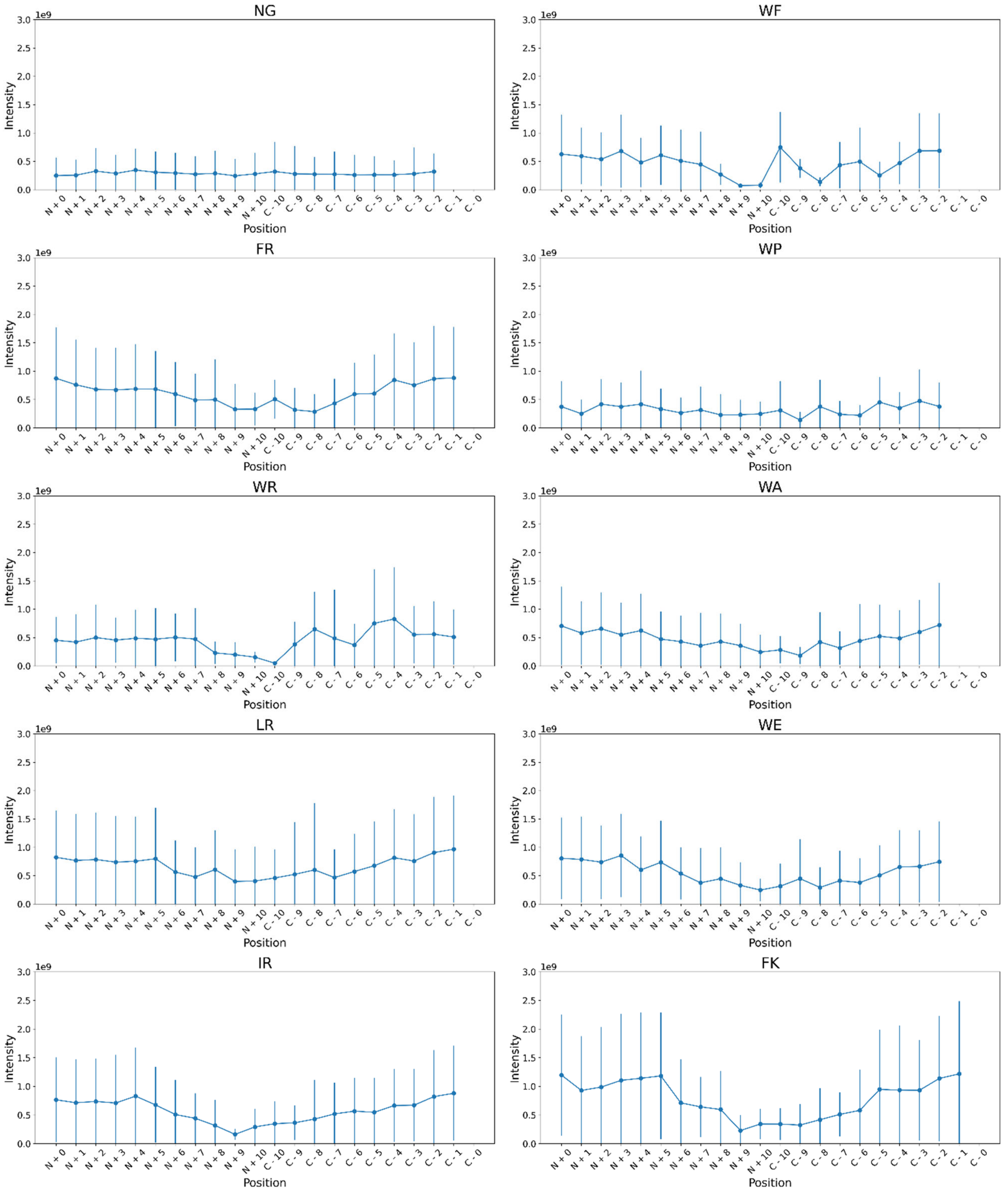


Figure S3: Mean intensity of top 10 dimer motifs from RM 3 based on position (N-terminus to N + 10 AAs and C – 10 AAs to C-terminus for the first AA in the dimer) within the peptide sequence. Plots are shown as position-dependent mean intensity with standard deviations within the filtered dataset.

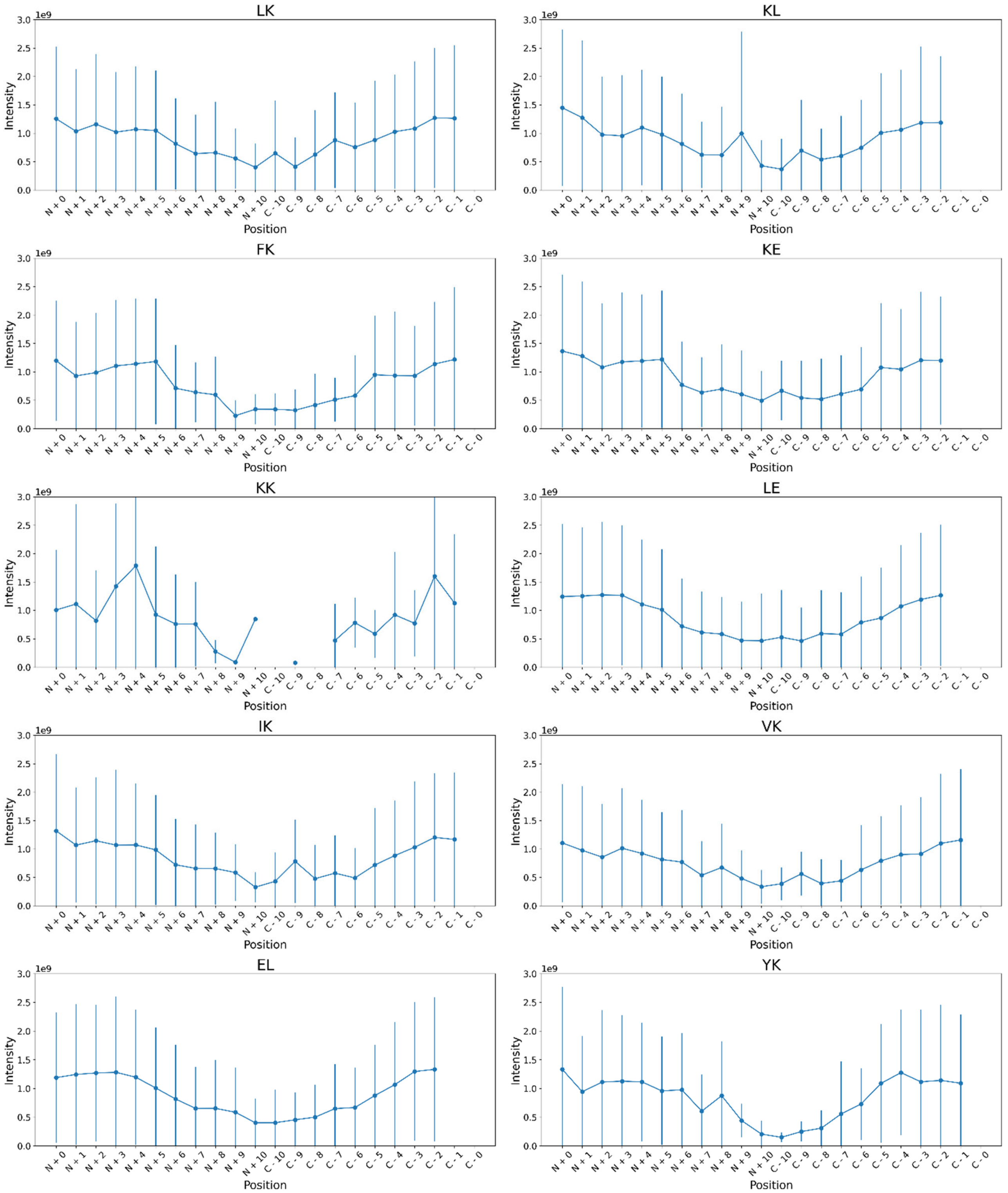


Figure S4: Mean intensity of top 10 dimer motifs from mean MS1 intensity data (Fig. 4) based on position (N-terminus to N + 10 AAs and C – 10 AAs to C-terminus for the first AA in the dimer) within the peptide sequence. Plots are shown as position-dependent mean intensity with standard deviations within the filtered dataset.

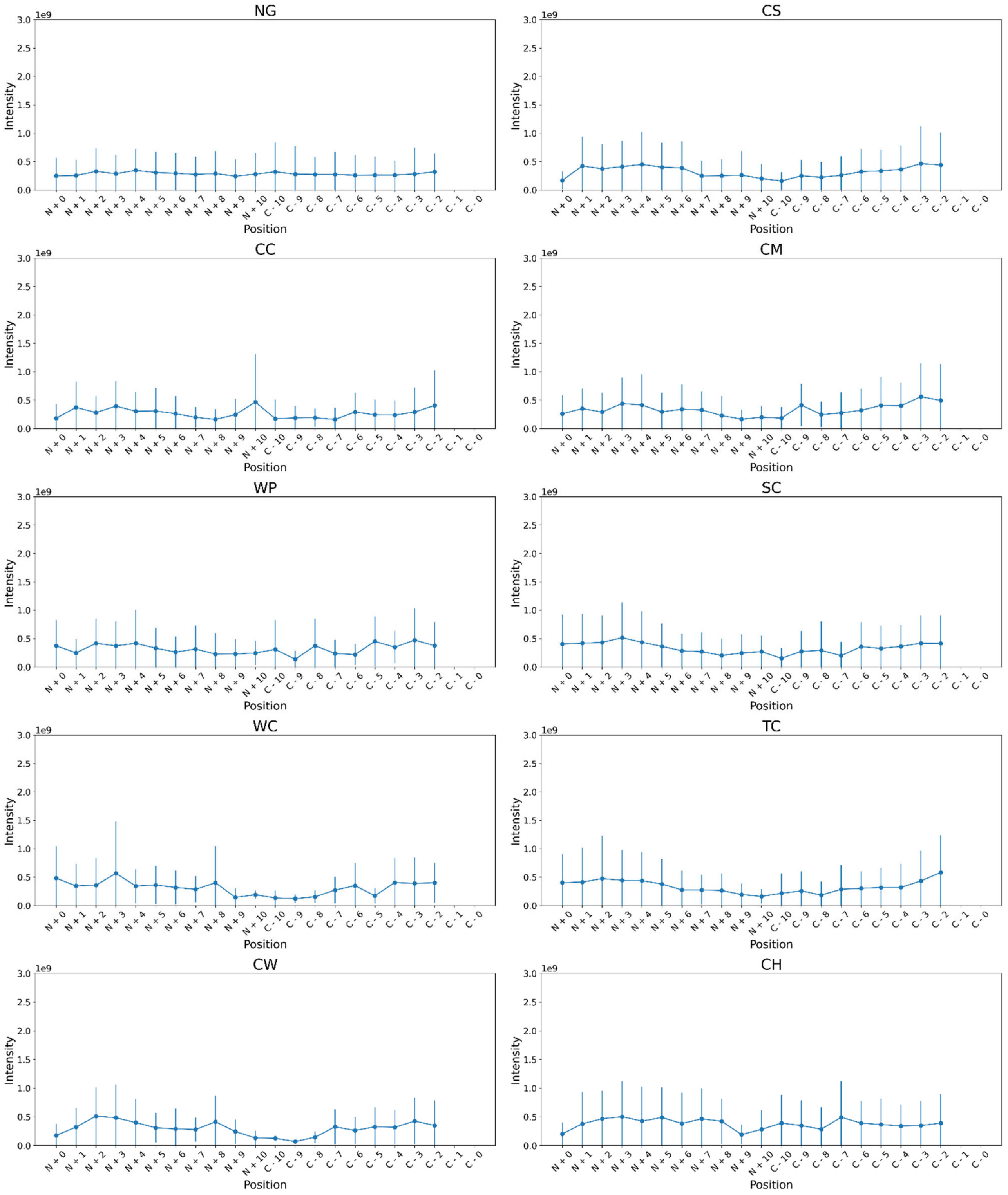


Figure S5: Mean intensity of bottom 10 dimer motifs from mean MS1 intensity data (Fig. 4) based on position (N-terminus to N + 10 AAs and C – 10 AAs to C-terminus for the first AA in the dimer) within the peptide sequence. Plots are shown as position-dependent mean intensity with standard deviations within the filtered dataset.
